# Gli3R-mediated inhibition of hedgehog signaling alters the embryonic transcriptome in zebrafish

**DOI:** 10.1101/2025.09.18.677168

**Authors:** Anna J. Moyer, Summer B. Thyme

**Affiliations:** Department of Biochemistry and Molecular Biotechnology, The University of Massachusetts Chan Medical School, Worcester, MA 01605, USA

**Keywords:** zebrafish, embryonic development, RNA-seq, hedgehog, Gli3R, Gli, ionocytes, ion homeostasis, Foxi3, H+-ATPase-rich

## Abstract

Hedgehog signaling is a conserved developmental pathway that patterns diverse tissues during vertebrate embryogenesis. In zebrafish, disruptions to the hedgehog pathway cause well-characterized defects in specific cell types including neurons and glia derived from the ventral neural tube. We inhibited hedgehog signaling by overexpressing the Gli3 repressor ubiquitously and performed bulk RNA-seq of 30 hours post-fertilization zebrafish embryos. Consistent with known roles of hedgehog signaling, we observed reduced expression of genes marking lateral floor plate, motor neurons, Kolmer-Agduhr cells, dopaminergic neurons, slow muscle cells, and anterior pituitary. Gene set enrichment analysis using marker genes derived from the Daniocell atlas also revealed downregulation of genes marking H+-ATPase-rich and Na+-K+-ATPase-rich ionocytes, which are located in the embryonic skin and are responsible for osmotic homeostasis. Reduced expression of ionocyte-specific transporter genes and the transcription factor *foxi3a* suggests that Gli activity may play a previously unrecognized role in the development of this cell type.

## INTRODUCTION

The hedgehog signaling pathway plays a critical role in vertebrate patterning (Ingham and McMahon 2001) and is also implicated in regeneration (Imokawa and Yoshizato 1997) and oncogenesis (Taipale et al. 2000). Depending on the context, hedgehog can act as a morphogen (Heemskerk and DiNardo 1994) or mitogen (Wechsler-Reya and Scott 1999) and specifies multiple distinct cell types across organ systems. In the canonical pathway, hedgehog ligands (Sonic, Indian, and Desert hedgehog) (Echelard et al. 1993) bind to the receptor Patched (Marigo et al. 1996; Stone et al. 1996), which relieves inhibition of the transducer Smoothened (Chen et al. 2001; Zhang et al. 2001). Activation of Smoothened shifts the balance between repressor and activator forms of the Gli transcription factors, which in turn modulates the transcription of target genes (Ruiz i Altaba 1999; Wang et al. 2000).

Experimental manipulation of hedgehog in the zebrafish model system has revealed essential roles in development (Hammerschmidt et al. 1996; Huang and Schier 2009; Shen et al. 2013). In the presumptive spinal cord, hedgehog secreted by the notochord and medial floor plate exposes the neighboring lateral floor plate (p3), pMN, and p0-p2 progenitor domains to a gradient of hedgehog levels, which contributes to specification of neural cell types including interneurons, motor neurons, Kolmer-Agduhr cells, oligodendrocytes, and astroglia [reviewed in (Danesin and Soula 2017)]. In addition to gross morphological phenotypes like cyclopia, ventral tail curvature, and rounded somites, inhibition of hedgehog in zebrafish can expand or reduce these ventral spinal cord progenitor domains and their derivative cells (Brand et al. 1996; Chen et al. 2001; Karlstrom et al. 1996).

However, subtle differences in phenotypes result from the specific method used to disrupt hedgehog signaling (i.e., mutations, morpholinos, overexpression of Gli repressors, and chemical inhibitors). These phenotypes may vary due to compensation between paralogs, incomplete or off-target effects of antagonists, maternal contribution of mRNA in homozygous mutants, and non-canonical or hedgehog ligand-independent Gli activity. For example, knockdown of *gli1* or *gli2b* in maternal-zygotic *smoothened* mutants reduces motor neuron number compared to mutants, suggesting that basal Gli activity contributes to motoneurogenesis (Mich and Chen 2011). Similarly, overexpression of the Gli3 repressor (Gli3R) in embryos treated with the Smoothened antagonist cyclopamine reduces the number of motor neurons compared to *smoothened* mutants (England et al. 2011). These experiments highlight how changes in the expression and processing of Gli can modify specification of cells in the ventral spinal cord even in the absence of upstream hedgehog signaling.

While previous work has manipulated Gli activity in targeted phenotypic assays, the transcriptome-wide consequences of Gli3R overexpression in zebrafish remain unexplored. To address this gap, we overexpressed a zebrafish *Gli3R* (*zGli3R*) transgene (Huang and Schier 2009) ubiquitously and performed bulk RNA sequencing (RNA-seq) of 30 hours post-fertilization (hpf) zebrafish embryos. Our findings suggest that Gli activity may be involved in the development of ionocytes. We anticipate that our dataset will be most relevant to zebrafish and developmental biologists who are interested in hedgehog signaling and cell-type specification during vertebrate development.

## MATERIALS & METHODS

### Zebrafish husbandry and transgenesis

Animal experiments were approved by the UMass Chan Institutional Animal Care and Use Committee (IACUC protocol 202300000053). Transgenic animals were generated in an Ekkwill (EK)-based wildtype strain, and both larvae and adult animals were maintained on a 14 hour/10 hour light/dark cycle at 28°C. The *zGli3R* transgene was subcloned from *pT2-hsp-zGli3R-EGFP* into Gateway *pDONR221* (Invitrogen) and then into a Gateway-compatible *Tol2* destination vector containing the *ubb* promoter (Mosimann et al. 2011) and the *myl7:GFP* transgenesis marker. The annotated plasmid map including full sequence is available in Supplementary File 1. To produce transgenic animals using *Tol2*, one-cell embryos were injected with 1 nL of 10 ng/µL *Tol2* mRNA (Kawakami et al. 2004) and 20 ng/µL plasmid. F0 embryos with expression of the *myl7:GFP* transgenesis marker were grown to adults.

### Sample collection

A mosaic F0 male was crossed to a wildtype female, and F1 embryos from one clutch were collected at 30 hpf for RNA-seq. Transgenic embryos and wildtype control siblings were sorted based on the *myl7:GFP* transgenesis marker. Four dechorionated embryos were pooled and frozen on dry ice for each biological replicate, and four biological replicates were collected per genotype. For live imaging, 2 days post-fertilization (dpf) and 3 dpf animals were anesthetized with MS-222 and live mounted in 2% low-melt agarose (Fisher Bioreagents, BP165-25) made in embryo medium with methylene blue. Images were obtained with a Leica M165 FC stereo microscope and K3C color camera.

### RNA-seq

RNA extraction, quality control, library preparation, and sequencing were conducted at Azenta Life Sciences (South Plainfield, NJ, USA) as follows: Total RNA was extracted from frozen tissue samples using Qiagen RNeasy Plus Universal mini kit following manufacturer’s instructions (Qiagen, Hilden, Germany). RNA samples were quantified using Qubit 3.0 Fluorometer (ThermoFisher Scientific, Waltham, MA, USA) and RNA integrity was checked with 4200 TapeStation (Agilent Technologies, Palo Alto, CA, USA).

RNA-seq libraries were prepared using the NEBNext Ultra II RNA Library Prep Kit for Illumina using manufacturer’s instructions (NEB, Ipswich, MA, USA). Briefly, mRNAs were initially enriched with Oligo d(T) beads. Enriched mRNAs were fragmented for 15 minutes at 94 °C. First strand and second strand cDNA were subsequently synthesized. cDNA fragments were end repaired and adenylated at 3′ ends, and universal adapters were ligated to cDNA fragments, followed by index addition and library enrichment by PCR with limited cycles. The sequencing library was validated on the Agilent TapeStation (Agilent Technologies, Palo Alto, CA, USA), and quantified by using Qubit 3.0 Fluorometer (Invitrogen, Carlsbad, CA) as well as by quantitative PCR (KAPA Biosystems, Wilmington, MA, USA).

The sequencing libraries were multiplexed and clustered onto a flowcell on the Illumina NovaSeq instrument according to manufacturer’s instructions. The samples were sequenced using a 2×150bp Paired End (PE) configuration. Image analysis and base calling were conducted by the NovaSeq Control Software (NCS). Raw sequence data (.bcl files) generated from Illumina NovaSeq was converted into fastq files and de-multiplexed using Illumina bcl2fastq 2.20 software. One mismatch was allowed for index sequence identification.

### RNA-seq analysis

Alignment and differential gene expression analysis of RNA-seq data were performed as previously described (Capps et al. 2025). Reads were aligned with the STAR aligner (2.7.3a-GCC 6.4.0-2.28) (Dobin et al. 2013) to GRCz11 release 104 using the Zebrafish Transcriptome Annotation version 4.3.2 (Lawson et al. 2020). Uniquely mapped reads ranged from 21.5-29.9M per sample. Transcripts with zero counts in four or more of eight total samples were removed, and raw counts were normalized using rlog counts in DESeq2 (Love et al. 2014). The default DESeq2 method was used for determining significance, and raw and adjusted (Benjamini-Hochberg) P values are in Supplementary Table 1. DESeq2 script is available on GitHub: https://github.com/thymelab/BulkRNASeq.

Downstream analysis of bulk RNA-seq data was conducted using R (Version 4.4.2). Gene set enrichment analysis (GSEA) was performed as previously described (Capps et al. 2025) using the ‘GSEA’ function from clusterProfiler (Version 4.14.6) (Wu et al. 2021). For chromosomal location GSEA, the gene set was generated using RefSeq Genes and Gene Predictions from the GRCz11 assembly, and genes were sorted by abs(log2 fold change) before GSEA. For Daniocell GSEA, the gene set was generated as previously described (Dang et al. 2025) and genes were sorted by log2 fold change before GSEA. Volcano plot, GSEA bar plot, and individual gene bar plots were created using ggplot2 (Version 3.5.2) (Wickham 2016). Heatmap was generated using the ‘heatmap.2’ function from gplots (Version 3.2.0) (Warnes et al. 2025). Code to generate figures is available in Supplementary File 2.

## RESULTS

### Differential gene expression in *Tg(ubb:zGli3R)* embryos

We cloned *zGli3R* downstream of the ubiquitin B (*ubb*) promoter and co-injected F0 embryos with *Tol2* to produce transgenic lines. Transgenic F1 offspring possessed severe morphological defects and died at larval stages (Fig. 1a). We isolated a mosaic F0 parent with high germline transmission and performed bulk RNA-seq on 30 hpf embryos pooled from a single clutch. Analysis of differentially expressed genes identified 888 upregulated genes and 1018 downregulated genes at a threshold of P adjusted (padj) <0.05 (Fig. 1b; Supplementary Table 1). Transgene insertion can cause enrichment of differentially expressed genes based on chromosomal location (White et al. 2022), and we observed a significant enrichment of genes on chromosome 4 (Supplementary Table 2).

**Fig. 1.**
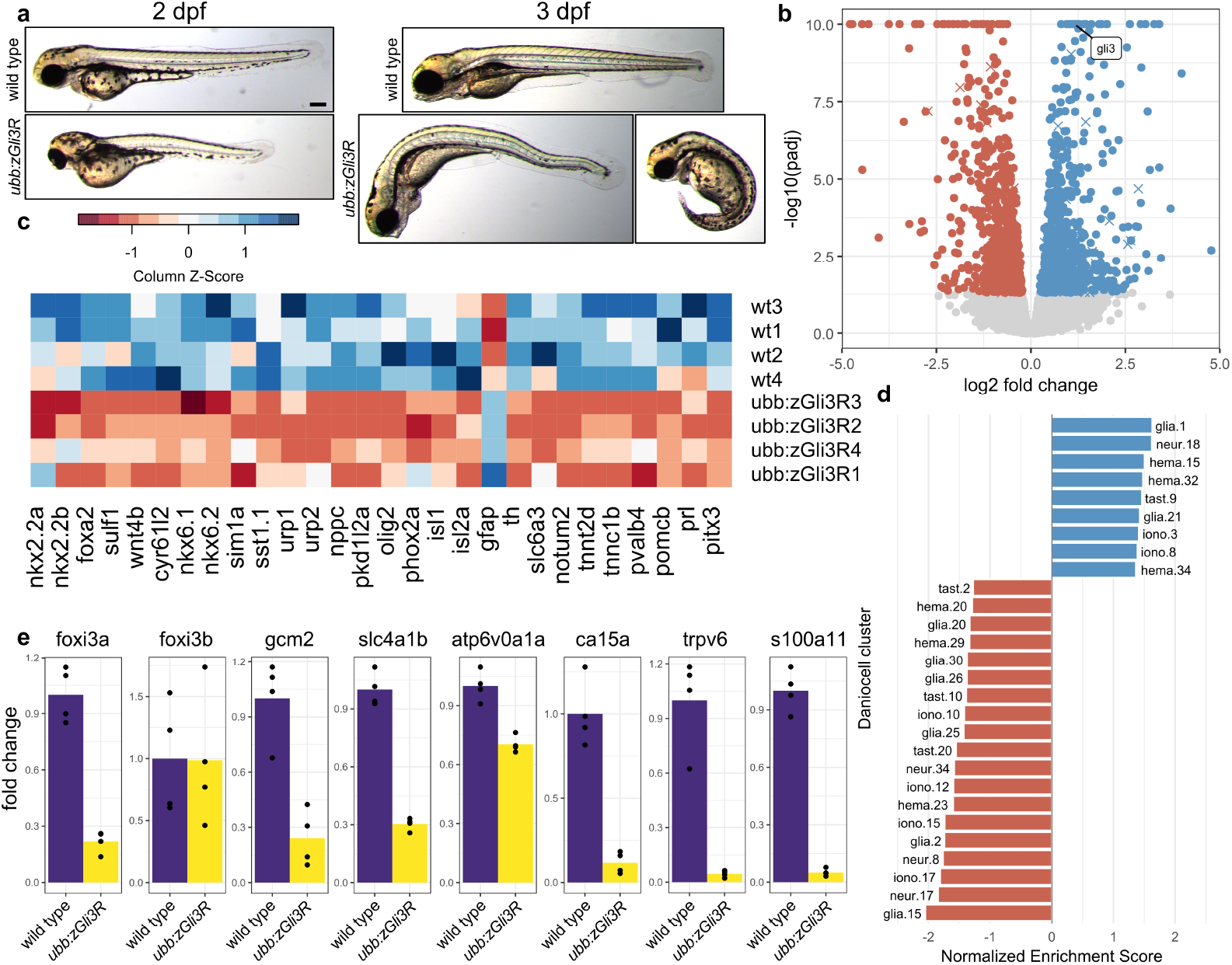
Morphological and transcriptomic phenotypes of *Tg(ubb:zGli3R)* embryos. a) Representative transgenic and wildtype animals at 2 dpf and 3 dpf. Scale bar = 200 μm. b) Volcano plot comparing gene expression in 30 hpf *Tg(ubb:zGli3R)* embryos and wildtype siblings. Genes with -log10(padj) greater than ten are plotted at ten. Transcripts encoded by chromosome 4 are denoted with X symbols and transcripts encoded by all other chromosomes are denoted with circles. n=4 replicates of 4 pooled embryos for transgenic and 4 replicates of 4 pooled embryos for wildtype siblings. c) Heat map displaying expression of selected marker genes. From left to right, genes mark progenitor domains and interneurons in the ventral spinal cord, Kolmer-Agduhr cells, motor neurons, radial glia/astrocytes, dopaminergic neurons, slow muscle cells, and anterior pituitary. d) Cell types identified with GSEA using gene sets from the Daniocell single-cell atlas (Sur et al. 2023). Red bars represent terms with a negative normalized enrichment score and blue bars represent terms with a positive score. e) Bar plots of normalized counts for genes expressed in ionocytes. P values and padj for all genes are available in Supplementary Table 1.

We next examined the expression of marker genes for cell types known to involve hedgehog signaling (Fig. 1c; Supplementary Table 1). Progenitor domains in the ventral spinal cord give rise to specific subtypes of neurons and glia, including interneurons, motor neurons, Kolmer-Agduhr cells, oligodendrocyte precursor cells (OPCs), and astrocytes [Reviewed in (Cucun et al. 2024; Danesin and Soula 2017)]. Genes marking most of these cell populations were downregulated or unchanged, but *gfap*, which is expressed in radial glia and astrocyte-like cells (Agius et al. 2004; Marcus and Easter 1995), was upregulated. Consistent with the literature, marker genes for dopaminergic neurons (Chen et al. 2001; Holzschuh et al. 2003), slow muscle cells (Blagden et al. 1997), and anterior pituitary (adenohypophysis) (Herzog et al. 2003) were also downregulated.

### Functional enrichment analysis of cell types using Daniocell single-cell atlas

GSEA using gene sets derived from single-cell expression data can reveal potential changes in cell composition (Capps et al. 2025; Dang et al. 2025; Moyer et al. 2025). GSEA with gene sets derived from the Daniocell single-cell atlas (Sur et al. 2023) highlighted 28 clusters with negative or positive enrichment compared to control (Fig. 1d; Supplementary Table 3). Several of the identified clusters with negative enrichment scores, including floor plate (glia.15 and glia.2), Kolmer-Agduhr cells (glia.30), spinal OPCs (glia.20 and glia.26), dopaminergic neurons of the diencephalon (neur.34), and anterior pituitary (tast.10) are consistent with the known roles of hedgehog during zebrafish development. Other cell types with negative enrichment scores included ventrolateral mesoderm (hema.20), hemangioblasts (hema.23), type III taste buds (tast.2), and multiple subtypes of H+-ATPase-rich ionocytes and ionocyte progenitors (iono.17, iono.15, iono.10, and iono.12). In contrast, olfactory sensory neurons (tast.9), lymphoid hematopoietic stem cells (hema.15), and primitive erythroblasts (hema.32) showed positive enrichment scores. Further investigation of known markers of ionocytes revealed downregulation of the master regulators *foxi3a* (Hsiao et al. 2007) and *gcm2* (Esaki et al. 2009; Shono et al. 2011) as well as genes expressed in H+-ATPase-rich ionocytes (*slc4a1b, atp6v0a1a, ca15a, ceacam1*, and *si:dkey-192d15*.*2*), and Na+-K+-ATPase-rich ionocytes (*trpv6* and *s100a11*) (Fig. 1e) (Sur et al. 2023).

## DISCUSSION

Although activation of hedgehog signaling removes Gli3R repressor activity in some contexts (Litingtung and Chiang 2000; Litingtung et al. 2002), *gli3* does not appear to be expressed before 12 hpf in zebrafish (England et al. 2011; Sur et al. 2023), and activator forms of Gli may possess basal levels of activity that are hedgehog ligand independent (Huang and Schier 2009; Karlstrom et al. 2003; Mich and Chen 2011). In this study, we explored the transcriptomic consequences of overexpressing *zGli3R* ubiquitously from the earliest stages of embryonic development. In contrast to experimental manipulations that inhibit hedgehog signaling, such as mutations in *smoothened* or cyclopamine treatment, ectopic expression of *zGli3R* has the potential to shift the balance of Gli activator and repressor forms even in the absence of hedgehog signaling.

We first examined marker genes for cell types previously shown to depend on hedgehog signaling (Fig. 1c) and then applied unbiased GSEA with Daniocell terms to discover new cell types that may be regulated by Gli (Fig. 1d). The validity of our experimental approach is supported by misexpression of known class II genes, which are activated by hedgehog signaling and include *nkx6*.*1, nkx6*.*2*, and *olig2* (Fig. 1c) (Briscoe et al. 2000; England et al. 2011; Guner and Karlstrom 2007). Genes marking Kolmer-Agduhr cells (*sst1*.*1, urp1*, and *urp2*), motor neurons (*phox2a* and *isl2a*), dopaminergic neurons (*th* and *slc6a3*), slow muscle cells (*notum2* and *tnnt2d*), and anterior pituitary (*pomcb*) were also significantly downregulated (Sur et al. 2023). GSEA using Daniocell gene sets identified several of the same populations, including anterior pituitary (tast.10), dopaminergic neurons (neur.34), and Kolmer-Agduhr cells (glia.30). However, GSEA also nominated cell types with less literature support for hedgehog involvement in zebrafish, including taste buds (tast.2), hemangioblasts (hema.23), and H+-ATPase-rich ionocytes (iono.17, iono.15, iono.10, and iono.12).

Ionocytes are specialized cells found in the embryonic skin that respond to osmotic changes in the aquatic environment and are functionally analogous to amniote renal tubular cells [Reviewed in (Hwang and Chou 2013)]. Progenitors derived from non-neural ectoderm give rise to both ionocytes and keratinocytes (Janicke et al. 2007). Expression of the transcription factors *foxi3a* and *foxi3b* in ionocyte progenitors activates Notch1 in neighboring cells, which in turn suppresses ionocyte cell fate and promotes keratinocyte development (Esaki et al. 2009; Hsiao et al. 2007; Janicke et al. 2007). The transcription factor *gcm2* also plays a role in the specification of ionocytes (Chang et al. 2009), and we observed reduced expression of *foxi3a* and *gcm2* but not of *foxi3b* (Fig. 1e).

Although hedgehog signaling has not been implicated in the specification of zebrafish ionocytes, it does regulate *gcm2* expression during development of the amniote parathyroid (Grevellec et al. 2011; Moore-Scott and Manley 2005). Moreover, the Daniocell single-cell atlas shows detectable expression of hedgehog components *smo, ptch2, gli1, gli2a, gli2b*, and *gli3* in non-neural ectoderm (neur.26). However, a recent study that used single-cell RNA-seq to track changes in cell abundance reported that cyclopamine treatment increases the abundance of non-keratinocyte epidermal progenitors and Na+-K+-ATPase-rich ionocytes (Barkan et al. 2025). Future hypothesis-based approaches will determine whether *zGli3R* overexpression affects ionocyte development via hedgehog signaling or by suppressing basal Gli activity.

## DATA AVAILABILITY STATEMENT

Plasmids are available upon request. Raw RNA-seq data and gene counts have been deposited in GEO (GSE307979). Processed data are available in Supplementary Table 1.

## ACKNOWLEDGEMENTS

The authors thank Dr. Peng Huang for providing the *pT2-hsp-zGli3R-EGFP* plasmid and research technician Vaishnavi Balaji for experimental support.

## Supplementary Materials

**Supplementary File 1:** Annotated plasmid map of *pTol2-ubb:zGli3R*.

**Supplementary File 2:** R code to reproduce figures.

**Supplementary Table 1:** DESeq2 output of differentially expressed genes.

**Supplementary Table 2:** GSEA by chromosomal location.

**Supplementary Table 3:** GSEA using terms derived from Daniocell atlas single-cell data.

