## Supplemental File 2 for "Gli3R-mediated inhibition of hedgehog signaling alters the embryonic transcriptome in zebrafish": supplementaryfile2_Rcode.html

zGli3R RNAseq


Code 

- Show All Code
- Hide All Code

### zGli3R RNAseq

###### Anna Moyer

#### *2025-09-16*

Make plots for RNAseq data collected from 30 hpf zebrafish embryos
ubb:zGli3r, line B.

### 1 Setup

#### 1.1 Set seed and working directory

```
#set random seed
set.seed(123)

#set working directory
setwd("C:/Users/annam/Desktop/lab work/bulkrnaseq/zgli3r/lineB")
```

#### 1.2 Load packages

```
library("tidyverse") 
library("DESeq2") #finding differentially expressed genes 
library("cowplot") #arranging plots into grids
library("viridis") #viridis color schemes
library("scales") #use to get color schemes for viridis
library("ggrepel") #repels text labels on plots
library("RColorBrewer") #pick colors
library("DT") #interactive and searchable tables of our GSEA results
library("GSEABase") #functions and methods for Gene Set Enrichment Analysis
library("Biobase") #base functions for bioconductor; required by GSEABase
library("GSVA") #Gene Set Variation Analysis, a non-parametric and unsupervised method for estimating variation of gene set enrichment across samples.
library("gprofiler2") #tools for accessing the GO enrichment results using g:Profiler web resources
library("clusterProfiler") # provides a suite of tools for functional enrichment analysis
library("msigdbr") # access to msigdb collections directly within R
library("enrichplot") # great for making the standard GSEA enrichment plots
library("ontologyIndex") #for parsing obo files
library("BaseSet") #for importing gaf file
library("plotly") #make interactive plots
library("lattice") #used for making manhattan plot
library("ggpubr") #calculating correlation coefficient 
library("colorspace") #colors for heatmap
library("ggraph") #library for making network graphs
library("svglite") #export svgs in loop
library("gplots") #make heatmaps
```

### 2 Chromosome analysis

#### 2.1 Add chromosome information

```
#import data frame with the previously calculated DEGs
lineB <- read.csv("input/zgli3r-lineB-homvswt_allresults_wt_hom-with-normalized.csv")[,2:9]

#add some information about pvalue
#limit padj to 1e-10
lineB <- lineB %>% mutate(padjbound = ifelse(lineB$padj < 1e-10, 1e-10, lineB$padj))

#limit pvalue to 1e-10
lineB <- lineB %>% mutate(pvaluebound = ifelse(lineB$pvalue < 1e-10, 1e-10, lineB$pvalue))

#import annotations for chromosome and TSS
chromosomeinfo <- readr::read_tsv(file="input/ncbi_refseqgenes")

#keep chromosome and gene name
chromosomeinfo <- chromosomeinfo %>% dplyr::select(chrom, name2, txStart)

#make empty columns to hold chromosome and txStart
lineB$chrom <- NA
lineB$txStart <- NA

#add chromosome and txStart information to data frame
for (x in 1:length(lineB$LLgeneAbbrev)) {
  gene <- lineB$LLgeneAbbrev[x]
  chromosomeinfosub <- chromosomeinfo %>% dplyr::filter(name2 == gene)
  chr <- chromosomeinfosub$chrom[1]
  txStart <- chromosomeinfosub$txStart[1]
  lineB$chrom[x] <- chr
  lineB$txStart[x] <- txStart
}

#add a column with chromosome number
lineB <- lineB %>% mutate(chromosomenumber = str_replace_all(chrom, c("chr1"="1", "chr2"="2", "chr3"="3", "chr4"="4", "chr5"="5", "chr6"="6", "chr7"="7", "chr8"="8", "chr9"="9", "chr10"="10", "chr11"="11", "chr12"="12", "chr13"="13", "chr14"="14", "chr15"="15", "chr16"="16", "chr17"="17", "chr18"="18", "chr19"="19", "chr20"="20", "chr21"="21", "chr22"="22", "chr23"="23", "chr24"="24", "chr25"="25")))
lineB <- distinct(lineB)

#export csv 
write.csv(lineB,file = "lineBDEG.csv")
```

### 3 Plotting DEGs by chromosome

Are differentially expressed genes located at a particular place in
the genome?

#### 3.1 Make Manhattan plot

This function is from: https://genome.sph.umich.edu/wiki/Code\_Sample:\_Generating\_Manhattan\_Plots\_in\_R

```
#import  comparisons (to skip running previous steps)
lineB <- read.csv("lineBdeg.csv")[,2:14]

#import  normalized counts 
lineBgenecounts <- read.csv("input/zgli3r-lineB-homvswt_normalized_reads_gene_list.csv")[,2:12]

#function for making manhattan plots
manhattan.plot<-function(chr, pos, pvalue, 
    sig.level=NA, annotate=NULL, ann.default=list(),
    should.thin=T, thin.pos.places=2, thin.logp.places=2, 
    xlab="Chromosome", ylab=expression(-log[10](p-value)),
    col=c("gray","darkgray"), panel.extra=NULL, pch=20, cex=0.8,...) {

    if (length(chr)==0) stop("chromosome vector is empty")
    if (length(pos)==0) stop("position vector is empty")
    if (length(pvalue)==0) stop("pvalue vector is empty")

    #make sure we have an ordered factor
    if(!is.ordered(chr)) {
        chr <- ordered(chr)
    } else {
        chr <- chr[,drop=T]
    }

    #make sure positions are in kbp
    if (any(pos>1e6)) pos<-pos/1e6;

    #calculate absolute genomic position
    #from relative chromosomal positions
    posmin <- tapply(pos,chr, min);
    posmax <- tapply(pos,chr, max);
    posshift <- head(c(0,cumsum(posmax)),-1);
    names(posshift) <- levels(chr)
    genpos <- pos + posshift[chr];
    getGenPos<-function(cchr, cpos) {
        p<-posshift[as.character(cchr)]+cpos
        return(p)
    }

    #parse annotations
    grp <- NULL
    ann.settings <- list()
    label.default<-list(x="peak",y="peak",adj=NULL, pos=3, offset=0.5, 
        col=NULL, fontface=NULL, fontsize=NULL, show=F)
    parse.label<-function(rawval, groupname) {
        r<-list(text=groupname)
        if(is.logical(rawval)) {
            if(!rawval) {r$show <- F}
        } else if (is.character(rawval) || is.expression(rawval)) {
            if(nchar(rawval)>=1) {
                r$text <- rawval
            }
        } else if (is.list(rawval)) {
            r <- modifyList(r, rawval)
        }
        return(r)
    }

    if(!is.null(annotate)) {
        if (is.list(annotate)) {
            grp <- annotate[[1]]
        } else {
            grp <- annotate
        } 
        if (!is.factor(grp)) {
            grp <- factor(grp)
        }
    } else {
        grp <- factor(rep(1, times=length(pvalue)))
    }
  
    ann.settings<-vector("list", length(levels(grp)))
    ann.settings[[1]]<-list(pch=pch, col=col, cex=cex, fill=col, label=label.default)

    if (length(ann.settings)>1) { 
        lcols<-trellis.par.get("superpose.symbol")$col 
        lfills<-trellis.par.get("superpose.symbol")$fill
        for(i in 2:length(levels(grp))) {
            ann.settings[[i]]<-list(pch=pch, 
                col=lcols[(i-2) %% length(lcols) +1 ], 
                fill=lfills[(i-2) %% length(lfills) +1 ], 
                cex=cex, label=label.default);
            ann.settings[[i]]$label$show <- T
        }
        names(ann.settings)<-levels(grp)
    }
    for(i in 1:length(ann.settings)) {
        if (i>1) {ann.settings[[i]] <- modifyList(ann.settings[[i]], ann.default)}
        ann.settings[[i]]$label <- modifyList(ann.settings[[i]]$label, 
            parse.label(ann.settings[[i]]$label, levels(grp)[i]))
    }
    if(is.list(annotate) && length(annotate)>1) {
        user.cols <- 2:length(annotate)
        ann.cols <- c()
        if(!is.null(names(annotate[-1])) && all(names(annotate[-1])!="")) {
            ann.cols<-match(names(annotate)[-1], names(ann.settings))
        } else {
            ann.cols<-user.cols-1
        }
        for(i in seq_along(user.cols)) {
            if(!is.null(annotate[[user.cols[i]]]$label)) {
                annotate[[user.cols[i]]]$label<-parse.label(annotate[[user.cols[i]]]$label, 
                    levels(grp)[ann.cols[i]])
            }
            ann.settings[[ann.cols[i]]]<-modifyList(ann.settings[[ann.cols[i]]], 
                annotate[[user.cols[i]]])
        }
    }
    rm(annotate)

    #reduce number of points plotted
    if(should.thin) {
        thinned <- unique(data.frame(
            logp=round(-log10(pvalue),thin.logp.places), 
            pos=round(genpos,thin.pos.places), 
            chr=chr,
            grp=grp)
        )
        logp <- thinned$logp
        genpos <- thinned$pos
        chr <- thinned$chr
        grp <- thinned$grp
        rm(thinned)
    } else {
        logp <- -log10(pvalue)
    }
    rm(pos, pvalue)
    gc()

    #custom axis to print chromosome names
    axis.chr <- function(side,...) {
        if(side=="bottom") {
            panel.axis(side=side, outside=T,
                at=((posmax+posmin)/2+posshift),
                labels=levels(chr), 
                ticks=F, rot=0,
                check.overlap=F
            )
        } else if (side=="top" || side=="right") {
            panel.axis(side=side, draw.labels=F, ticks=F);
        }
        else {
            axis.default(side=side,...);
        }
     }

    #make sure the y-lim covers the range (plus a bit more to look nice)
    prepanel.chr<-function(x,y,...) { 
        A<-list();
        maxy<-ceiling(max(y, ifelse(!is.na(sig.level), -log10(sig.level), 0)))+.5;
        A$ylim=c(0,maxy);
        A;
    }

    xyplot(logp~genpos, chr=chr, groups=grp,
        axis=axis.chr, ann.settings=ann.settings, 
        prepanel=prepanel.chr, scales=list(axs="i"),
        panel=function(x, y, ..., getgenpos) {
            if(!is.na(sig.level)) {
                #add significance line (if requested)
                panel.abline(h=-log10(sig.level), lty=2);
            }
            panel.superpose(x, y, ..., getgenpos=getgenpos);
            if(!is.null(panel.extra)) {
                panel.extra(x,y, getgenpos, ...)
            }
        },
        panel.groups = function(x,y,..., subscripts, group.number) {
            A<-list(...)
            #allow for different annotation settings
            gs <- ann.settings[[group.number]]
            A$col.symbol <- gs$col[(as.numeric(chr[subscripts])-1) %% length(gs$col) + 1]    
            A$cex <- gs$cex[(as.numeric(chr[subscripts])-1) %% length(gs$cex) + 1]
            A$pch <- gs$pch[(as.numeric(chr[subscripts])-1) %% length(gs$pch) + 1]
            A$fill <- gs$fill[(as.numeric(chr[subscripts])-1) %% length(gs$fill) + 1]
            A$x <- x
            A$y <- y
            do.call("panel.xyplot", A)
            #draw labels (if requested)
            if(gs$label$show) {
                gt<-gs$label
                names(gt)[which(names(gt)=="text")]<-"labels"
                gt$show<-NULL
                if(is.character(gt$x) | is.character(gt$y)) {
                    peak = which.max(y)
                    center = mean(range(x))
                    if (is.character(gt$x)) {
                        if(gt$x=="peak") {gt$x<-x[peak]}
                        if(gt$x=="center") {gt$x<-center}
                    }
                    if (is.character(gt$y)) {
                        if(gt$y=="peak") {gt$y<-y[peak]}
                    }
                }
                if(is.list(gt$x)) {
                    gt$x<-A$getgenpos(gt$x[[1]],gt$x[[2]])
                }
                do.call("panel.text", gt)
            }
        },
        xlab=xlab, ylab=ylab, 
        panel.extra=panel.extra, getgenpos=getGenPos, ...
    );
}

#select columns to include in manhattan plot
myTopHits.df <- lineB %>% dplyr::select(chromosomenumber, txStart, pvaluebound)

#filter to only include chromosomes 1-25
myTopHits.df <- myTopHits.df %>% dplyr::filter(chromosomenumber %in% 1:25)

#make chromosomes plotted in order
myTopHits.df$chromosomenumber <- factor(myTopHits.df$chromosomenumber, levels = 1:25)

#omit NA values
myTopHits.df <- na.omit(myTopHits.df)

#make colors 
manhattancol <- c(replicate(1, c("lightgrey", "darkgrey")), "lightgrey", "#21908CFF", replicate(10, c("lightgrey", "darkgrey")))
  
#can filter at this step if desired
manhattan <- myTopHits.df 

#make manhattan plot
manhattan.plot(manhattan$chromosomenumber, manhattan$txStart, manhattan$pvaluebound, should.thin=F, col=manhattancol)
```

```
#export manhattan plots at 800x500 pixels
```

Next will test whether these genes are formally overrepresented using
GSEA.

#### 3.2 Make custom GSEA annotation with zebrafish genes and chromosomes

```
#import annotations for chromosome and gene name
chromosomeinfo <- readr::read_tsv(file="input/ncbi_refseqgenes")

#keep chromosome and gene name
chromosomeinfo <- chromosomeinfo %>% dplyr::select(chrom, name2)

#change column names
colnames(chromosomeinfo) <- c("chromosome", "genename")

#select only chromosomes 1-25
chromosomeinfo <- chromosomeinfo %>% dplyr::filter(chromosome %in% paste("chr", 1:25, sep=""))

#keep only distinct rows
chromosomeinfo <- distinct(chromosomeinfo)
```

#### 3.3 Check for enrichment of one chromosome in DEGs

```
# Perform GSEA using clusterProfiler
#remove duplicates
duplicategenes <- dplyr::filter(lineB %>%
  distinct() %>%
  group_by(LLgeneAbbrev) %>%
  dplyr::count(), n !=1)$LLgeneAbbrev

#remove duplicate rows from GSEAgenes
mydata.df.sub <- lineB %>% dplyr::filter(!LLgeneAbbrev %in% duplicategenes)

#keep only the columns we need for GSEA
mydata.df.sub <- dplyr::select(mydata.df.sub, LLgeneAbbrev, log2FoldChange)

#sort by abs(foldchange)
mydata.df.sub$log2FoldChange <- abs(mydata.df.sub$log2FoldChange)

# construct a named vector
mydata.gsea <- mydata.df.sub$log2FoldChange
names(mydata.gsea) <- as.character(mydata.df.sub$LLgeneAbbrev)
mydata.gsea <- sort(mydata.gsea, decreasing = TRUE)

# run GSEA using the 'GSEA' function from clusterProfiler
myGSEA.res <- GSEA(mydata.gsea, TERM2GENE=chromosomeinfo, verbose=FALSE, scoreType="pos", maxGSSize=2000, pvalueCutoff = 1)
myGSEA.df <- as_tibble(myGSEA.res@result)

# view results as an interactive table
datatable(myGSEA.df, 
          extensions = c('KeyTable', "FixedHeader"), 
          options = list(keys = TRUE, searchHighlight = TRUE, pageLength = 10, lengthMenu = c("10", "25", "50", "100"))) %>%
  formatRound(columns=c(2:10), digits=2)

#export
#write.csv(myGSEA.df, "chromosomeGSEA.csv")
```

Genes on chromosome 4 are enriched in line B.

### 4 Analysis of DEGs

```
#count upregulated and downregulated genes
#upregulated genes
dim(lineB %>% dplyr::filter(padj < 0.05 & log2FoldChange > 0))
```

```
## [1] 888  13
```

```
#There are 888 genes with padj <0.05 and positive fold change

#downregulated genes
dim(lineB %>% dplyr::filter(padj < 0.05 & log2FoldChange < 0))
```

```
## [1] 1018   13
```

```
#There are 1018 genes with padj <0.05 and negative fold change

#make a copy of the data
myTopHits.df <- lineB

#add column about whether chr is 4
myTopHits.df <- myTopHits.df %>% dplyr::mutate(chrom = replace_na(chrom, "none"))
myTopHits.df <- myTopHits.df %>% mutate(chromosome = ifelse(chrom == "chr4", "chromosome 4", "other chromosome"))

#add a column about genes that are upregulated
myTopHits.upregulated <- myTopHits.df %>% dplyr::filter(padj < 0.05 & log2FoldChange > 0)
myTopHits.upregulated$category <- "upregulated"

#add a column about genes that are downregulated
myTopHits.downregulated <- myTopHits.df %>% dplyr::filter(padj < 0.05 & log2FoldChange < 0)
myTopHits.downregulated$category <- "downregulated"

#add a column to color gli3
myTopHits.gli3 <- myTopHits.df %>% dplyr::filter(LLgeneAbbrev == "gli3")
myTopHits.gli3$category <- "gli3"

#genes to color
myTopHits.labels <- bind_rows(myTopHits.upregulated, myTopHits.downregulated, myTopHits.gli3)

#change order that points are plotted
myTopHits.labels <- myTopHits.labels %>% arrange(match(category,  c("gli3", "upregulated", "downregulated")), desc(category))

genestolabel <- c("gli3")

#subset to only include labeled genes
myTopHits.labels.all <- myTopHits.df %>% dplyr::filter(LLgeneAbbrev %in% genestolabel)

#make all points other
myTopHits.df <- myTopHits.df %>% dplyr::mutate(mutation = "other")

#make the plot
lineB_volcano <- ggplot() +
  geom_point(data=myTopHits.df, aes(y=-log10(padjbound), x=log2FoldChange, shape = chromosome, color = mutation), size=2) +
  geom_point(data=myTopHits.labels, aes(y=-log10(padjbound), x = log2FoldChange, color=category, shape=chromosome),  size=2, show.legend = T) +
  theme_bw() +
  coord_cartesian(xlim = c(-5, 5), ylim = c(-0.5, 10.5), expand = FALSE) +
  ylab("-log10(padj)") + 
  xlab("log2 fold change") +
  geom_label_repel(data=myTopHits.labels.all, aes(x=log2FoldChange, y=-log10(padjbound), label=LLgeneAbbrev), force = 2, nudge_y = -1, size = 2.5, max.overlaps = Inf, show.legend = FALSE, color = "black") + #label selected genes
  theme_bw() +
  scale_color_manual(values = c("#c96351", "black", "lightgray", "#5993c2"), name = "pathway") +
  scale_shape_manual(values=c(4, 16)) 

lineB_volcano
```

```
#export 525x325
```

#### 4.1 Heatmap

```
mySelectedGenes <- c("nkx2.2a", "nkx2.2b", "foxa2", "sulf1", "wnt4b", "cyr61l2", "nkx6.1", "nkx6.2", "sim1a",  "sst1.1", "urp1", "urp2", "nppc", "pkd1l2a", "olig2", "phox2a", "isl1", "isl2a", "gfap", "th", "slc6a3", "notum2", "tnnt2d", "tnnc1b", "pvalb4", "pomcb", "prl", "pitx3")

#filter replicate data with list of selected genes
mySelectedGenes_exp <- lineBgenecounts %>% filter(LLgeneAbbrev %in% mySelectedGenes)

#reorder data table based on myselectedgenes
mySelectedGenes_exp <- mySelectedGenes_exp[match(mySelectedGenes, mySelectedGenes_exp$LLgeneAbbrev),]

#turn replicate data into a matrix
mySelectedGenes.matrix <- as.matrix(mySelectedGenes_exp[,4:11])

#add row names (genes) to data matrix
rownames(mySelectedGenes.matrix) <- mySelectedGenes_exp$LLgeneAbbrev

#mySelectedGenes.matrix <- mySelectedGenes.matrix[!rowSums(is.na(mySelectedGenes.matrix)),]
#mySelectedGenes.matrix <- mySelectedGenes.matrix[rowSums(mySelectedGenes.matrix[])>0,]

#you may (or may not) want to cluster your selected genes
hr <- hclust(as.dist(1-cor(t(mySelectedGenes.matrix), method="pearson")), method="complete") #cluster rows by pearson correlation
hc <- hclust(as.dist(1-cor(mySelectedGenes.matrix, method="spearman")), method="average") #cluster columns by spearman correlation

#get color
myheatcolors3 <- brewer.pal(name="RdBu", n=11)

#make heatmap
heatmap.2(mySelectedGenes.matrix, 
          Rowv=NA, 
          Colv=as.dendrogram(hc), 
          col=myheatcolors3, scale="row", 
          density.info="none", trace="none", dendrogram = "none")
```

```
#make horizontal
matrix_t <- t(mySelectedGenes.matrix)

#you may (or may not) want to cluster your selected genes
hr_sample <- hclust(as.dist(1-cor(t(mySelectedGenes.matrix), method="pearson")), method="complete") #cluster rows by pearson correlation
hc_gene <- hclust(as.dist(1-cor(mySelectedGenes.matrix, method="spearman")), method="average") #cluster columns by spearman correlation


#make heatmap
heatmap.2(matrix_t, 
          Rowv=as.dendrogram(hc_gene), 
          Colv=NA, 
          col=myheatcolors3, scale="column", 
          density.info="none", trace="none", dendrogram = "none", cexCol=1.5, margins = c(8, 4))
```

```
#export 900x400
```

#### 4.2 GSEA with Daniocell

```
singlecellmarkers <- read.csv("input/daniocell_GSEAterms.csv")

singlecellmarkers <- dplyr::select(singlecellmarkers, cluster, gene)

#remove duplicates
duplicategenes <- dplyr::filter(lineB %>%
  distinct() %>%
  group_by(LLgeneAbbrev) %>%
  dplyr::count(), n !=1)$LLgeneAbbrev

#remove duplicate rows from GSEAgenes
GSEAgenes <- lineB %>% dplyr::filter(!LLgeneAbbrev %in% duplicategenes)

# pull out data that we need
mydata.df.sub <- dplyr::select(GSEAgenes, LLgeneAbbrev, log2FoldChange)

#get rid of duplicates
mydata.df.sub <- mydata.df.sub %>% dplyr::filter(LLgeneAbbrev %in% mydata.df.sub$LLgeneAbbrev[!duplicated(mydata.df.sub$LLgeneAbbrev)])

# construct a named vector
mydata.gsea <- mydata.df.sub$log2FoldChange
names(mydata.gsea) <- as.character(mydata.df.sub$LLgeneAbbrev)
mydata.gsea <- sort(mydata.gsea, decreasing = TRUE)

# run GSEA with daniocell markers
myGSEA.res <- GSEA(mydata.gsea, TERM2GENE=singlecellmarkers, verbose=FALSE, seed=TRUE)

#convert to DF
myGSEA.df <- as_tibble(myGSEA.res)

#look at top terms
datatable(myGSEA.df, 
          extensions = c('KeyTable', "FixedHeader"), 
          options = list(keys = TRUE, searchHighlight = TRUE, pageLength = 10, lengthMenu = c("10", "25", "50", "100"))) %>%
  formatRound(columns=c(2:10), digits=2)
```

```
#export
#write.csv(myGSEA.df, "lineB-GSEA-daniocell.csv")
```

#### 4.3 Make bar plot from line B GSEA

```
#select columns of GSEA to plot
GSEAbardata <- dplyr::select(myGSEA.df, ID, NES)

# Color based on value
color <- ifelse(GSEAbardata$NES < 0, "#c96351", "#5993c2")

#make plot
barplot <- ggplot(GSEAbardata, aes(x = reorder(ID, NES), y = NES)) +
  geom_bar(stat = "identity",
           show.legend = FALSE,
           fill = color,     
           color = "white") +
  geom_hline(yintercept = 0, color = 1, lwd = 0.2) +
  geom_text(aes(label = ID,
                hjust = ifelse(NES < 0, 1.1, -0.1),
                vjust = 0.5), size = 3) +
  xlab("Daniocell cluster") +
  ylab("Normalized Enrichment Score") +
  scale_y_continuous(limits = c(-2.5, 2.5)) +
  coord_flip() +
  theme_minimal() +
  theme(axis.text.y = element_blank(),  # Remove Y-axis texts
        axis.ticks.y = element_blank(), # Remove Y-axis ticks
        panel.grid.major.y = element_blank()) # Remove horizontal grid

barplot
```

```
#export 335 x 500 pixels
```

#### 4.4 Make bar plots for selected genes

```
#pick out a genes to plot
geneofinterest <- c("foxi3a", "foxi3b", "slc4a1b", "trpv6", "s100a11", "atp6v0a1a", "gcm2", "ca15a") 
#geneofinterest <- c("ceacam1")

#make new columns with mean counts of wt
genecountslogfc <- lineBgenecounts
genecountslogfc$wtavg <- rowMeans(genecountslogfc[,c(5,7,9,11)])

#divide columns by mean counts of wt
genecountslogfc <- genecountslogfc %>% mutate(across(colnames(genecountslogfc)[4:11], function(x) x/wtavg))

#pull out data for a select gene
generepdata <- genecountslogfc %>% dplyr::filter(LLgeneAbbrev %in% geneofinterest)

#get rid of unnecessary columns
generepdata <- generepdata[,c(1,4:11)]

#pivot longer to make tidy
generepdata <- generepdata %>% pivot_longer(!LLgeneAbbrev, names_to = "sample", values_to = "foldchange")

#add condition information based on sampleName
generepdata$genotype <- generepdata$sample

#substitute condition name based on replicate
generepdata <- generepdata %>% mutate(genotype = str_replace_all(genotype, c("^het.*"="het", "^hom.*"="hom", "^wt.*" = "wt")))

#make genotype a factor
generepdata$genotype <- factor(generepdata$genotype, levels = c("wt", "het", "hom"))

#make plots manually split by age
colors <- c("#472D7BFF", "#FDE725FF", "#21908CFF") #set colors

#make an empty list
plot_list = list()

#make all the plots
for (z in 1:length(geneofinterest)) {
  genedata <- generepdata %>% dplyr::filter(LLgeneAbbrev == geneofinterest[z]) 
  plot1 <- ggplot(genedata, aes(fill=genotype, y=foldchange, x=genotype)) + 
    geom_bar(position="dodge", stat="summary", width=1) +
    geom_point(position = position_dodge(width = .9)) +
    labs(title = genedata$LLgeneAbbrev[z]) +
    ylab("fold change") +
    xlab("") + 
    theme_bw() +
    scale_fill_manual(values = colors) +
    scale_x_discrete(labels=c('wild type', 'ubb:zGli3R')) +
    theme(legend.position="none", axis.text.x = element_text(angle = 45, vjust = 1, hjust = 1), plot.title = element_text(hjust = 0.5))
  plot_list[[z]] <- plot1
}

plot_list
```

```
## [[1]]
```

```
## 
## [[2]]
```

```
## 
## [[3]]
```

```
## 
## [[4]]
```

```
## 
## [[5]]
```

```
## 
## [[6]]
```

```
## 
## [[7]]
```

```
## 
## [[8]]
```

```
# Save plots to svg Makes a separate file for each plot.
for (i in 1:length(geneofinterest)) {
    file_name = paste("lineB_foldchange", geneofinterest[i], ".svg", sep="")
    svglite(file_name, width=1.5, height=4)
    print(plot_list[[i]])
    dev.off()
}
```

### 5 Session info

Packages and versions necessary to reproduce the results in this
report.

```
sessionInfo()
```

```
## R version 4.4.2 (2024-10-31 ucrt)
## Platform: x86_64-w64-mingw32/x64
## Running under: Windows 10 x64 (build 19045)
## 
## Matrix products: default
## 
## 
## locale:
## [1] LC_COLLATE=English_United States.utf8 
## [2] LC_CTYPE=English_United States.utf8   
## [3] LC_MONETARY=English_United States.utf8
## [4] LC_NUMERIC=C                          
## [5] LC_TIME=English_United States.utf8    
## 
## time zone: America/New_York
## tzcode source: internal
## 
## attached base packages:
## [1] stats4    stats     graphics  grDevices utils     datasets  methods  
## [8] base     
## 
## other attached packages:
##  [1] gplots_3.2.0                svglite_2.2.1              
##  [3] ggraph_2.2.1                colorspace_2.1-1           
##  [5] ggpubr_0.6.0                lattice_0.22-6             
##  [7] plotly_4.10.4               BaseSet_1.0.0              
##  [9] ontologyIndex_2.12          enrichplot_1.26.6          
## [11] msigdbr_24.1.0              clusterProfiler_4.14.6     
## [13] gprofiler2_0.2.3            GSVA_2.0.7                 
## [15] GSEABase_1.68.0             graph_1.84.1               
## [17] annotate_1.84.0             XML_3.99-0.18              
## [19] AnnotationDbi_1.68.0        DT_0.33                    
## [21] RColorBrewer_1.1-3          ggrepel_0.9.6              
## [23] scales_1.4.0                viridis_0.6.5              
## [25] viridisLite_0.4.2           cowplot_1.1.3              
## [27] DESeq2_1.46.0               SummarizedExperiment_1.36.0
## [29] Biobase_2.66.0              MatrixGenerics_1.18.1      
## [31] matrixStats_1.5.0           GenomicRanges_1.58.0       
## [33] GenomeInfoDb_1.42.3         IRanges_2.40.1             
## [35] S4Vectors_0.44.0            BiocGenerics_0.52.0        
## [37] lubridate_1.9.4             forcats_1.0.0              
## [39] stringr_1.5.1               dplyr_1.1.4                
## [41] purrr_1.0.4                 readr_2.1.5                
## [43] tidyr_1.3.1                 tibble_3.3.0               
## [45] ggplot2_3.5.2               tidyverse_2.0.0            
## [47] knitr_1.49                  tinytex_0.56               
## [49] rmarkdown_2.29             
## 
## loaded via a namespace (and not attached):
##   [1] splines_4.4.2               bitops_1.0-9               
##   [3] ggplotify_0.1.2             R.oo_1.27.1                
##   [5] polyclip_1.10-7             lifecycle_1.0.4            
##   [7] rstatix_0.7.2               vroom_1.6.5                
##   [9] MASS_7.3-61                 crosstalk_1.2.1            
##  [11] backports_1.5.0             magrittr_2.0.3             
##  [13] sass_0.4.10                 jquerylib_0.1.4            
##  [15] yaml_2.3.10                 ggtangle_0.0.6             
##  [17] DBI_1.2.3                   abind_1.4-8                
##  [19] zlibbioc_1.52.0             R.utils_2.13.0             
##  [21] yulab.utils_0.2.0           tweenr_2.0.3               
##  [23] GenomeInfoDbData_1.2.13     irlba_2.3.5.1              
##  [25] tidytree_0.4.6              codetools_0.2-20           
##  [27] DelayedArray_0.32.0         ggforce_0.4.2              
##  [29] DOSE_4.0.1                  tidyselect_1.2.1           
##  [31] aplot_0.2.6                 UCSC.utils_1.2.0           
##  [33] farver_2.1.2                ScaledMatrix_1.14.0        
##  [35] jsonlite_2.0.0              tidygraph_1.3.1            
##  [37] Formula_1.2-5               systemfonts_1.2.3          
##  [39] tools_4.4.2                 treeio_1.30.0              
##  [41] snow_0.4-4                  Rcpp_1.0.14                
##  [43] glue_1.8.0                  gridExtra_2.3              
##  [45] SparseArray_1.6.2           xfun_0.51                  
##  [47] qvalue_2.38.0               HDF5Array_1.34.0           
##  [49] withr_3.0.2                 fastmap_1.2.0              
##  [51] rhdf5filters_1.18.1         caTools_1.18.3             
##  [53] digest_0.6.37               rsvd_1.0.5                 
##  [55] timechange_0.3.0            R6_2.6.1                   
##  [57] gridGraphics_0.5-1          textshaping_1.0.1          
##  [59] GO.db_3.20.0                gtools_3.9.5               
##  [61] dichromat_2.0-0.1           RSQLite_2.4.1              
##  [63] R.methodsS3_1.8.2           generics_0.1.4             
##  [65] data.table_1.17.4           graphlayouts_1.2.2         
##  [67] httr_1.4.7                  htmlwidgets_1.6.4          
##  [69] S4Arrays_1.6.0              pkgconfig_2.0.3            
##  [71] gtable_0.3.6                blob_1.2.4                 
##  [73] SingleCellExperiment_1.28.1 XVector_0.46.0             
##  [75] htmltools_0.5.8.1           carData_3.0-5              
##  [77] fgsea_1.32.4                png_0.1-8                  
##  [79] SpatialExperiment_1.16.0    ggfun_0.1.8                
##  [81] rstudioapi_0.17.1           tzdb_0.5.0                 
##  [83] reshape2_1.4.4              rjson_0.2.23               
##  [85] nlme_3.1-166                curl_6.3.0                 
##  [87] cachem_1.1.0                rhdf5_2.50.2               
##  [89] KernSmooth_2.23-24          parallel_4.4.2             
##  [91] pillar_1.10.2               grid_4.4.2                 
##  [93] vctrs_0.6.5                 car_3.1-3                  
##  [95] BiocSingular_1.22.0         beachmat_2.22.0            
##  [97] xtable_1.8-4                evaluate_1.0.3             
##  [99] magick_2.8.7                cli_3.6.4                  
## [101] locfit_1.5-9.12             compiler_4.4.2             
## [103] rlang_1.1.5                 crayon_1.5.3               
## [105] ggsignif_0.6.4              labeling_0.4.3             
## [107] plyr_1.8.9                  fs_1.6.6                   
## [109] stringi_1.8.7               BiocParallel_1.40.0        
## [111] assertthat_0.2.1            babelgene_22.9             
## [113] Biostrings_2.74.1           lazyeval_0.2.2             
## [115] GOSemSim_2.32.0             Matrix_1.7-1               
## [117] hms_1.1.3                   patchwork_1.3.0            
## [119] sparseMatrixStats_1.18.0    bit64_4.6.0-1              
## [121] Rhdf5lib_1.28.0             KEGGREST_1.46.0            
## [123] igraph_2.1.4                broom_1.0.8                
## [125] memoise_2.0.1               bslib_0.9.0                
## [127] ggtree_3.14.0               fastmatch_1.1-6            
## [129] bit_4.6.0                   ape_5.8-1                  
## [131] gson_0.1.0
```
